# eDNA replicates, polymerase and amplicon size impact inference of richness across habitats

**DOI:** 10.1101/2024.11.19.624235

**Authors:** Jarl Andreas Anmarkrud, Fabricio dos Anjos Santa Rosa, Lisbeth Thorbek, Audun Schrøder-Nielsen, Silvana Melo Sviggum, Jonathan Stuart Ready, Hugo J. de Boer, Quentin Mauvisseau

**Affiliations:** Natural History Museum, University of Oslo, Norway; Group for Integrated Biological Investigation, Center for Advanced Studies of Biodiversity, Federal University of Pará, Belém, Brazil

## Abstract

Environmental DNA-based monitoring has been increasingly used in the last decade to monitor biodiversity in aquatic and terrestrial systems. Molecular-based surveys now allow quick and reliable production of baseline knowledge of species community composition on a large scale, allowing better understanding of ecosystem function and mitigation of stressors linked to anthropogenic activities. Despite this, technical hurdles often remain, and the impact of replicates, PCR polymerases and amplicon size on the recovered species richness is still poorly understood. Here, we conducted a large controlled experiment, with bulk samples collected from terrestrial, marine and freshwater environments to assess the impact of natural and technical replicates, PCR polymerases with different degrees of fidelity or proofreading activity, as well as amplicon size on species richness recovery across habitats. In this study, we consistently found variations in sample species richness depending on PCR polymerase choice. We further demonstrate the dissimilar impacts between natural and technical replicates on species richness recovery, and the necessity of increasing natural replications in eDNA based surveys. We highlight the benefits and limitations of replication strategies, polymerase choice and amplicon size across terrestrial, marine and freshwater habitats, and provide recommendations to increase the reliability of future eDNA-based metabarcoding studies.

## Introduction

In the last decade, environmental DNA (eDNA) metabarcoding has become an increasingly applied method for assessing biodiversity in a variety of substrates and ecological communities (Ruppert et al., 2019; Blackman et al., 2024). eDNA refers to both organismal and extra organismal DNA isolated from the environment, and can be present as (i) dissolved DNA, (ii) organelle, (iii) cell or (iv) DNA bound to suspended particles (Mauvisseau et al., 2022; Rodriguez-Ezpeleta et al., 2021). Detection of these traces in soil, water, sediment and even air has revolutionised the monitoring capacity for ecological studies and biodiversity assessments worldwide (Goldberg et al., 2018; Beermann et al., 2020; Bohmann & Lynggaard, 2022a; Carvalho et al., 2024). Initial studies investigated the efficiency and reliability of this monitoring method compared to traditional techniques (Shaw et al., 2016; Watts et al., 2019; Hallam et al., 2021; Basset et al., 2022; Keck et al., 2022) and morphology analyses (Berry et al., 2015; Emilson et al., 2017; Schenk et al., 2020; Brantschen et al., 2021), and showed that eDNA could be used to detect biodiversity from environmental substrates that were difficult or impossible to study using morphology, e.g., dietary analysis from faeces, vegetation reconstruction from ancient sedimentary DNA (Alsos et al., 2016; Guillerault et al., 2017; Alsos et al., 2018; Ribas et al., 2021; Queiroz et al., 2024; Rosa et al., 2024). Indeed, compared to traditional taxonomic biodiversity screening, DNA metabarcoding is more sensitive, yields more comprehensive taxonomic datasets, is more cost-efficient and relies less on taxonomic expertise (Ji et al., 2013; Fediajevaite et al., 2021). While the method is not likely to replace botanists making floristic inventories, lepidopterists catching butterflies or ornithologists spotting birds, it is likely to find increasing adoption and application in assessment and monitoring scenarios where it excels compared to traditional methods (Gogarten et al., 2019; Mas-Carrió et al., 2022; Milla et al., 2022). Coupled with the involvement of citizen science allowing habitat monitoring on a large scale, reanalysis of old samples or museum collection it offer a promising avenue for future ecological and biodiversity mapping studies (Bi et al., 2013; Burian et al., 2023; Jeunen et al., 2024).

DNA extracted from environmental samples consists of DNA molecules shed from multiple organisms, many of them present with relatively few molecules, and multiple natural and technical replicates are needed to obtain reliable results and decrease the risks of false negatives (Ficetola et al., 2015, 2016; Erickson et al., 2019; Mauvisseau et al., 2019; Burian et al., 2021; Darling et al., 2021; Acharya-Patel et al., 2024). eDNA sampling protocols vary across environments and studies, and optimal design often differs depending on research questions and cost-efficiency (Smart et al., 2016; Lugg et al., 2018; Wilcox et al., 2018). In the eDNA metabarcoding analytical workflow, DNA extraction and potential inhibitor removal (McKee et al., 2014; Djurhuus et al., 2017; Tsuji et al., 2019; Uchii et al., 2019; Pawlowski et al., 2021) is followed by PCR amplification of the targeted genetic markers. Metabarcoding then relies on the analysis of a relatively short targeted genetic region, a “barcode”, amplified by polymerase chain reaction (PCR) and sequenced using high-throughput sequencing (Valentini et al., 2009; Thomsen & Willerslev, 2015; Taberlet et al., 2018).

PCR amplification is highly prone to bias, and significant stochasticity is observed in results (Zinger et al., 2019; Bohmann et al., 2021). This is exacerbated if the DNA concentration and purity of the eDNA template is low (Acharya-Patel et al., 2024). As the driver of the PCR reaction, the polymerase enzyme plays a critical role in the amplification of DNA extracted in environmental samples and can be associated with biases depending on primers used and environments analysed (Acharya-Patel et al., 2024). Indeed, primers are used to amplify DNA from targeted organisms or groups prior to NGS sequencing, and primer choices might vary depending on the sampled environments or study design, as they will have a great impact on the recovered richness (Freeland, 2016; Hajibabaei et al., 2019; Shu et al., 2021; Espinosa Prieto et al., 2024). The resulting sequence data, “the barcodes”, are thereafter mapped to publicly available genetic databases for identification of ZOTUs (i.e. Zero Radius Operational Taxonomic Units, also called Amplicon Sequence Variants -ASVs) or OTUs (Operational Taxonomic Units, also called Molecular Operational Taxonomic Units - MOTUs) (Antich et al., 2021). ZOTUs are generated through denoising, and refer to all correct biological sequences, while OTUs are generated through clustering and refer to subsets of the correct biological sequences (Edgar, 2010, 2013, 2016), but both serve as a molecular proxy to identify a taxon. However, prior to the generation of ZOTUs or OTUs, the DNA metabarcoding workflow consists of several steps and strategic experimental decisions that may all influence the outcome and composition of molecular proxies (Deiner et al., 2017).

Due to these known biases, careful experimental design is necessary to overcome potential limitations (Rees et al., 2015; Zaiko et al., 2018; Kestel et al., 2022; Couton et al., 2023). To mitigate such hurdles, we designed a large controlled experiment to investigate the impacts of both natural and technical replicate numbers, polymerases, and primers (targeting different fragment sizes and genes) on freshwater, marine and soil bulk samples (Figure 1). While some of these parameters have been studied individually previously, the current study is the first one investigating their combined effect on such a large scale. Here, we tested if (i) polymerases with low fidelity performed better than polymerases with high fidelity to recover richness, (ii) if natural and technical replicates had similar or different impacts on richness recovery, (iii) if the combination of all these variables lead to similar results across marine, freshwater and terrestrial habitats, and (iv) if these effects were consistent using primer sets amplifying shorter and larger DNA fragments targeting different mitochondrial genes.

**Figure 1:**
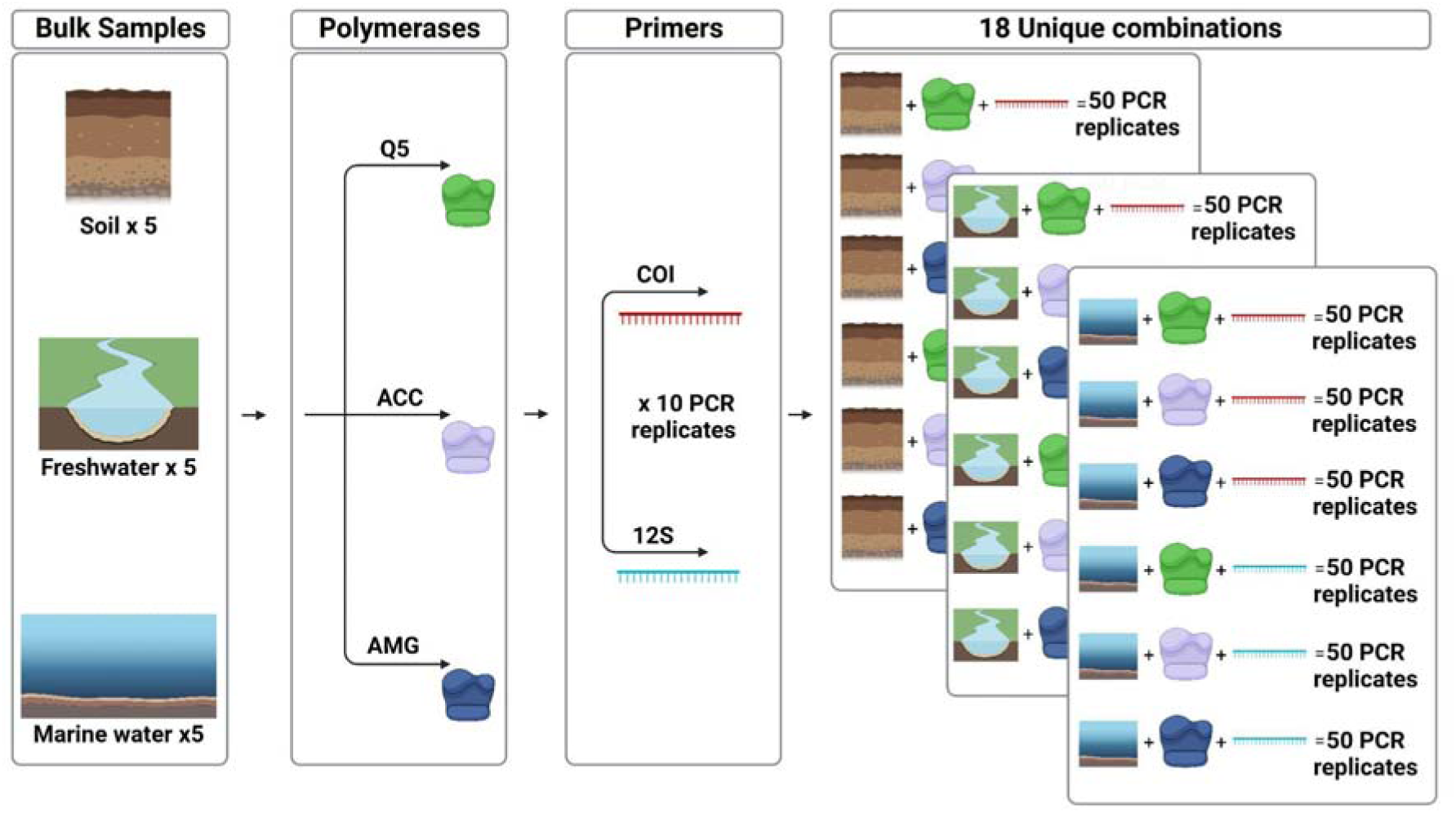
Study design. Bulk Samples from soil, marine and freshwater environments were created, and DNA was extracted from five natural replicates of each bulk sample. Then, for each environment, a combination of three polymerases and two markers were analysed, with ten PCR replicates conducted for each natural replicate. This led to the analysis of 18 Unique Combinations across the workflow of our analysis, with 50 PCR replicates per combination for each environment, and a total of 60 negative controls additionally included in the analysis. The figure was created using BioRender.

## Methods

### Sample collection and extraction

#### Freshwater

Freshwater was collected from the surface of the Drengsrudbekken river (59.8346N, 10.43014E), Asker, Norway, and from the surface of the Akerselva river (59.9166N, 10.7615E), Oslo, Norway on February 1^st^ 2023. No environmental parameters were collected at these sampling locations. Equal amounts of water from both rivers were pooled and thoroughly mixed together in a 15 L sterile jerrycan to create a freshwater artificial mock community. Then, a total of five independent subsamples (referred to later as natural replicates) of 1 L each were filtered through 0.8 μm Whatman (Cytiva, Germany) Cellulose Nitrate membrane filters (25 mm diameter) using sterile 25 mm Swinnex (Merck Millipore, Germany) Filter Holders and a Vampire Sampler (Bürkle GmbH, Germany) system. Environmental DNA was extracted from the filters using the DNeasy Blood & Tissue Kit (Qiagen) with slight modification (volumes of ATL, proteinase K, AL buffers and ethanol were doubled, and the lysis step was conducted overnight).

#### Marine

Marine water was collected from 5 locations across a transect in the Oslo fjord (59.6603N, 10.6051E; 59.6606N, 10.6100E; 59.6610N, 10.6144E; 59.6627N, 10.6181E; 59.6633N, 10.6230E) on January 18^th^ 2023. An equal amount of water from each sampling location (i.e. 2 L), was pooled and thoroughly mixed together in a 15 L sterile jerrycan to create a marine artificial mock community. Following this, five independent subsamples (i.e. natural replicates) of 1 L each were filtered through 0.8 μm Whatman (Cytiva, Germany) Cellulose Nitrate membrane filters (25 mm diameter) using sterile 25 mm Swinnex (Merck Millipore, Germany) Filter Holders and a Vampire Sampler (Bürkle GmbH, Germany) system. Environmental DNA was extracted from the filters using the same kit and method described earlier regarding the freshwater eDNA samples.

#### Terrestrial

A total of 5 soil samples were collected in Vesterøy, Hvaler, Norway in Summer 2018 (2 samples in 59.0906N; 10.8994E, one sample in 59.08667N; 10.86798E and 2 samples in 59.0900N; 10.8648E) (see full sample description in Ariza et al. 2024). At each location, various debris (i.e. stones, rocks, living or dead plant parts) were removed to expose the upper organic soil layer. Then, a 50 ml falcon tube was inserted into the soil top layer until it was entirely filled, and capped before being stored at -20 Ill back in the laboratory. From each of these 5 initial soil samples, 3 g were merged (15g total) in a 100 ml grinding chamber (IKA-Werke GmbH & Co) and homogenised using a IKA Tube mill 100 (IKA-Werke GmbH & Co) to create an artificial soil mock community. Following this, five independent subsamples of 250 mg each were extracted using the Qiagen PowerSoil Kit (Qiagen) following protocol with slight modifications (for the lysis step, a Mini-Beadbeater (Techtum) was used to homogenise the samples for 2 minutes at 25 Hz).

To ensure the absence of contamination, one negative control (i.e. sample without any biological contents) was also included in each of the extraction workflows of the respective marine, freshwater and terrestrial samples. DNA concentration of all DNA extracts was measured using a high sensitivity kit on a Qubit 2.0 spectrophotometer (Invitrogen) to assess the extraction success. All DNA samples were then diluted to a concentration of 1-2 ng/µl before conducting PCR amplification.

#### PCR amplification

Amplification targeted a short fragment (approx 106 bp) of the mitochondrial 12S rRNA using the 12S-V5F 5’-ACTGGGATTAGATACCCC-3’ and 12S-V5R 5’-TAGAACAGGCTCCTCTAG-3’ primers (Riaz et al., 2011; Kelly et al., 2014), and another longer fragment (approx 313 bp) of the mitochondrial COI region using the mlCOIintF 5’-GGWACWGGWTGAACWGTWTAYCCYCC-3’ and Fol-degen-rev 5’-TANACYTCNGGRTGNCCRAARAAYCA-3’ primers (Yu et al., 2012; Leray et al., 2013). PCRs were conducted using three different polymerase mastermixes: (i) Q5 High-Fidelity 2X Master Mix (New England Biolabs), a mastermix high fidelity polymerase reported with >280X higher fidelity than Taq polymerase (Potapov & Ong, 2017), referred to as “Q5” throughout the manuscript, (ii) AmpliTaq Gold 360 mastermix (ThermoFisher), a mastermix with a Taq polymerase designed for high sensitivity and requiring low copy number of target for amplification (Taberlet et al., 2018), and is referred to as “AMG’’ throughout the manuscript and (iii) Accustart II PCR Toughmix (QuantaBio), a mastermix designed for degraded and/or challenging material with a high degree of known PCR inhibitors, and referred to as “ACC” throughout the manuscript. The Q5 polymerase used in this study does not require heat activation and, in contrast to AMG and ACC (who are both hotstart enzymes), has a proofreading activity.

#### PCR optimization

Prior to metabarcoding amplification, amplification success for each primer set was assessed through a gradient analysis on a subset of samples. In brief, PCRs were conducted in a 15 µl final volume, including 1X of the polymerase mastermix, 0.3 µM of each primer and 1.5 µl of template DNA diluted to a concentration of 1-2 ng/µl. It should be noted that both Q5 and AMG mastermixes were provided with a GC enhancer, an additive which prohibits formation of secondary structures. As a result, PCR reactions were performed in parallel with and without the GC enhancer for the Q5 and AMG mastermixes in order to verify if this reagent improved amplification quality. These initial tests allowed us to identify the annealing temperature for each primer set and polymerase generating the highest amplicon yield without producing extra fragments outside of the target size range. Amplification success was verified using 1.2% agarose gel electrophoresis, and visualised using the ImageLab v6.0 software on a GelDoc XR+ system (BioRad). Based on this initial testing, the PCR were run as follows: Q5 and Accustart: 1X enzyme mastermix, 0.3 µM primers, 1.5 µl of diluted DNA template and nuclease free water to a final volume of 15 µl. Similar conditions were used for the Amplitaq Gold master mix except we also included 20% (v/v) of the provided GC enhancer. Details regarding the final PCRs conditions for each primer set and polymerase can be found in Supplementary Table 1.

None of the extraction negative controls (i.e. samples without any biological contents) yielded amplicons during the initial tests aiming to optimise the PCR reactions for each primers and polymerases combination. As a result, these extraction negative controls were not included for amplifications alongside the soil, freshwater and marine eDNA samples. However, a total of 60 NTC or non-template control (PCR amplification where the DNA template was replaced with ddH_2_O) were included in the amplification workflow (see Supplementary Table S2). All PCR amplifications were performed using indexed primers following the dual-index design as in Fadrosh et al., (2014). All indexed primers used in this study were ordered without modification and using desalt purification.

#### Metabarcoding analysis

All unique eDNA samples analysed in this study were amplified using ten PCR replicates (see Figure 1). Together with the analysis of 18 unique combinations (i.e. three habitats, three polymerases, two amplicon sizes) with 50 PCR replicates per combination (i.e. five natural replicates analysed each with 10 PCR replicates), this resulted in the analysis of 960 unique amplicons (900 eDNA samples and 60 NTCs). All amplicons were visualised through a 1.2% agarose gel electrophoresis and relative quantities of each amplicon were estimated using the software ImageLab v6.0 (BioRad). Equimolar amounts of each amplicon were merged into separate pools based on marker and enzyme used in amplification using a Biomek4000 liquid handling robot (Beckman Coulter). The pools were cleaned using 10% of Illustra ExoProStar (Cytiva) and AMPure XP Beads (Beckman Coulter) (0.8X ratio COI, 1.2X ratio 12S). Quality control of the cleaned amplicon pools were performed on a Fragment Analyzer system (Agilent). Because of the different fragment lengths of the two markers, we sequenced amplicons from each primer set on two separate Illumina MiSeq flow cells (COI: MiSeq V3 300 PE, 12S: MiSeq V2 250 PE) .

#### Bioinformatics

Bioinformatics data processing, filtering and cleaning was performed as in (Carvalho et al., 2024; Carvalho et al., 2024) with slight modification depending on the amplicon’s libraries. In brief, forward and reverse raw sequencing reads were merged using PEAR 0.9.3 (Zhang et al., 2014) and demultiplexing was performed using the *ngsfilter* algorithm from OBITools (Boyer et al., 2015). The *obigrep* algorithm from OBITools was used to remove fragments >420 bp from the COI library, and >200 bp from the amplicons of the 12S library. Additional filtering was conducted using the USEARCH algorithm (Edgar, 2010), and sequences <100 bp were removed from both 12S and COI amplicon libraries, as well as sequences showing less than 10 occurrences in the datasets. Finally, 12S and COI datasets were denoised using the UNOISE algorithm (Edgar, 2016) to retrieve ZOTUs and clustered using the UPARSE algorithm (Edgar, 2013) to retrieve OTUs, both using the default 97% identity threshold in USEARCH (Edgar, 2010, 2013, 2016). Both clustering and denoising were performed to retrieve subsets of the correct biological sequences (OTUs) as well as all correct biological sequences (ZOTUs), to assess later the effect of polymerases and proofreading activities on OTU/ZOTU richness. As we created artificial mock communities to assess the impacts of replication, polymerase and amplicon size on eDNA samples from various habitats, this manuscript mainly focuses on raw OTU/ZOTU richness (i.e. without taxonomic assignment). However, taxonomic assignment and similar downstream analysis were additionally performed and can be found in (Figures S1, S2, S3, S4, Table S3, File S1). We performed strict filtering to remove any potential false positive detection across the 12S and COI datasets. We first investigated whether each ZOTU/OTU was detected in any of the 60 negative controls (NTCs). If any reads were observed for a given ZOTU/OTU in the negative controls, we then retained the highest number reads for this ZOTU/OTU in the negative controls, and subtracted this number from all PCR replicates from all samples across the dataset. This conservative approach was performed for all ZOTUs/OTUs across both 12S and COI datasets. This was done to ensure the absence of contamination or other sequencing errors.

#### Statistical analysis

Statistical analyses and data visualisation were performed using R software (R Foundation for Statistical Computing, 2021). Comparison between OTU/ZOTU richness obtained for each combination was done using the Kruskal-Wallis test from the *stats* package (version 4.4.1), followed by Dunn’s post hoc test using the *dunn.test* package (version 1.3.6). The Kruskal-Wallis test is a non-parametric method used to assess statistically significant differences between the medians of three or more independent groups, in this case, the three types of DNA polymerase: ACC, AMG, and Q5. After identifying an overall difference, Dunn’s test was applied as a post hoc analysis to perform pairwise comparisons, determining which specific polymerase pairs showed significant differences in OTU/ZOTU richness (Harper et al., 2023). A Bonferroni correction was used to adjust for multiple comparisons, reducing the chances of type I errors by dividing the significance threshold by the number of comparisons (Steinke et al., 2022). A significance level of 0.05 was used for both tests. To visually represent the OTU/ZOTU richness across polymerases, a violin plot was created using the *ggplot2* package (version 3.5.1) (Wickham, 2016), with significant p-values highlighted on the plot to emphasise differences between polymerase groups. Violin plots provide a clear visualisation of richness values by showing the data distribution and density, alongside key statistical measures such as centralization and dispersion, thus facilitating the detection of any unusual patterns or extreme values (Casals & Daunis-i-Estadella, 2023). Rarefaction curves investigating the impacts of natural and technical replicates for each combination were generated using the “specaccum” function from the *vegan* package (version 2.6.8). The number of natural replicates ranged from one to five, with richness values being aggregated accordingly. For each natural replicate the 10 technical replicates were processed for each DNA polymerase combination. These rarefaction curves were then plotted using the *ggplot2* package (version 3.5.1) (Wickham, 2016), visualising richness across the technical replicates for each polymerase combination.

## Results

A total of 14 297 880 and 10 283 473 raw reads were obtained following metabarcoding amplification and sequencing of the 12S and COI libraries. Following bioinformatics processes including data cleaning and denoising, as well as removing singletons and ZOTUs showing less than 10 reads across the datasets, a total of 8 286 609 reads for 541 unique ZOTUs and 5 424 606 reads for 13 143 unique ZOTUs remained for both 12S and COI datasets respectively. The clustering approach resulted in a total of 8 306 467 reads for 403 unique OTUs and 5 390 422 reads for 9 063 unique OTUs remaining for both 12S and COI datasets respectively, removing singletons and OTUs showing less than 10 reads across the COI and 12S datasets. Following strict additional filtering of the datasets using negative controls to remove potential false positive detection, a total of 6 941 654 reads with 541 unique ZOTUs remained for the 12S dataset, and a total of 5 191 140 reads with 13 143 unique ZOTUs remained for the COI dataset. In comparison, the OTU production pipeline resulted in a total of 6 953 229 reads with 403 unique OTUs for the 12S dataset, and a total of 5 145 307 reads with 9 063 unique OTUs for the COI dataset. A higher number of OTUs can be expected, as ZOTUs would generate clusters with low read counts which would be filtered out when removing unique reads below 10 singletons. Additional information regarding retained ZOTUs and OTUs for both COI and 12S datasets following taxonomic assignment can be found in (Figures S1, S2). OTUs and ZOTUs tables can be found in (Tables S4-S11).

Out of the 900 unique amplicons (i.e. 450 amplicons per primer set after the exclusion of negative controls), amplification success varied across polymerases, with ACC showing higher read numbers across all environments using the COI primer set, but with more similar read numbers across all polymerases and environments using 12S primer set (Figure S3). This was consistent across both the OTU and ZOTU approaches. We found that Q5 retrieved significantly lower OTU and ZOTU richness than ACC and AMG for large COI amplicons (*p-values* ≤0.001) across all environments. We found a similar effect with shorter 12S amplicons using the ZOTU approach in the soil environment (*p-values* ≤0.05). A significant difference was also found between Q5 and AMG using the OTU approach in the soil environment (*p-values* ≤0.05). However, no significant differences were found between ACC and AMG for both short 12S amplicons (*p-value* =0.57) and larger COI amplicons (*p-value* =0.99) (Figure 2). Due to the inherent variation of species composition across marine, freshwater and terrestrial environments, and primer efficiency (i.e. primer sets designed to target different organisms, genes, and DNA fragment sizes) we did not compare the species richness retrieved across habitats and markers. However, details regarding significant differences across polymerases per habitat and per primer sets can be found in Table S3, for both OTU and ZOTU approaches.

**Figure 2:**
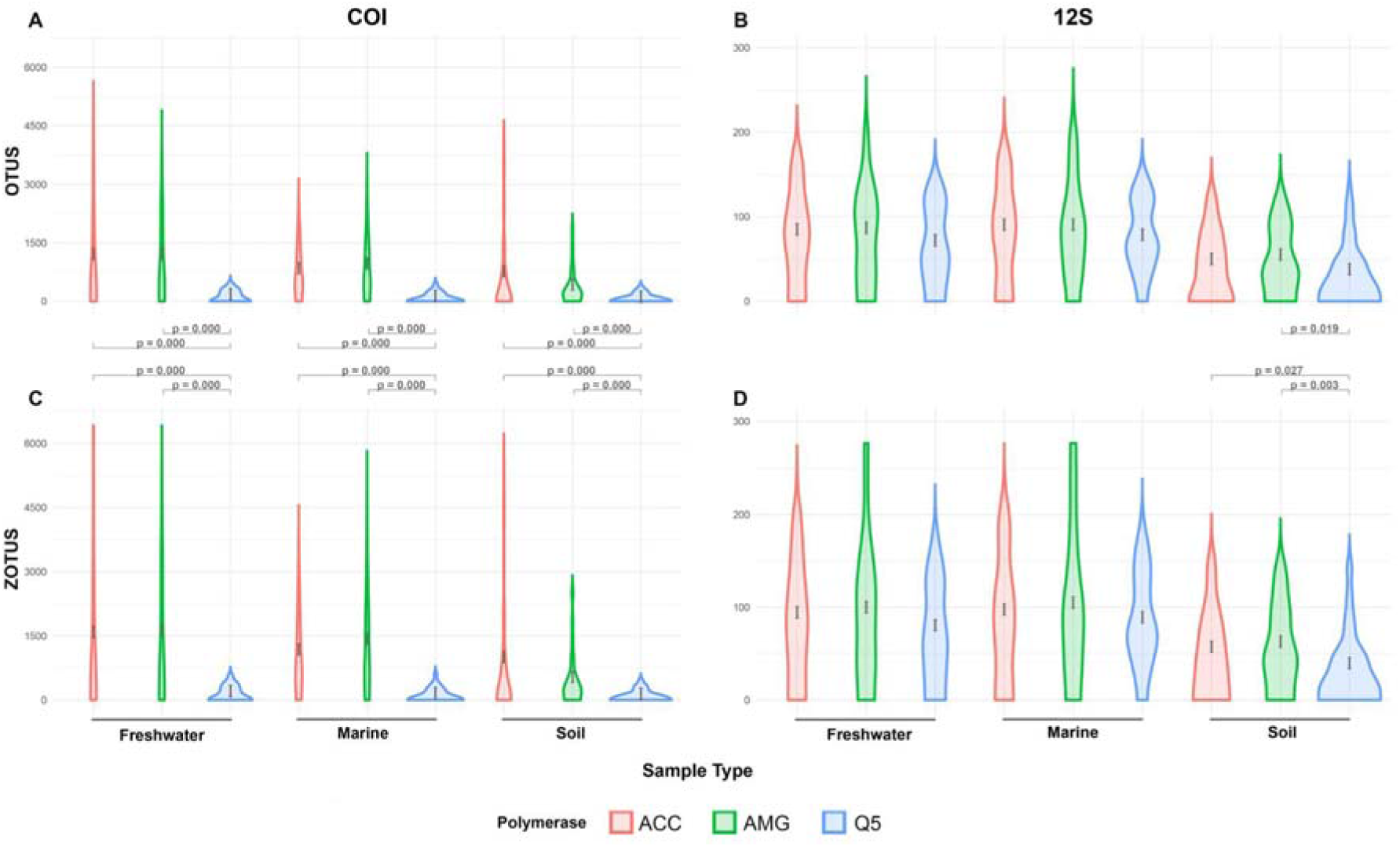
Recovered richness visualised in violin plots using non-linear Box-Cox transformation for each three polymerases showed that Q5 retrieved significantly lower OTU and ZOTU richness than ACC and AMG for large COI amplicons (*p-values* ≤0.001) across all environments. A similar effect was found with shorter 12S amplicons using the ZOTU approach in the soil environment (*p-values* ≤0.05). Significant difference was also found between Q5 and AMG using the OTU approach in the soil environment (*p-values* ≤0.05). The figure highlights: (A) COI amplicons analysed using the OTU approach with significant p-values; (B) 12S amplicons generated under the OTUs approach; (C) COI amplicons analysed using the ZOTU approach; and (D) 12S amplicons generated under the ZOTU approach. The only approach where no significant results were found was the 12S amplicons using the ZOTU approach.

Finally, we found that across all tested combinations (i.e. three polymerases across three habitats), the number of natural replicates (i.e. number of filter or soil samples) had a dissimilar impact compared to the number of technical replicates (i.e. PCR replicate) on the recovery of OTU and ZOTU richnesses. The increase of natural replicates allowed a greater recovery of OTU and ZOTU richnesses than the increase of technical replicates (Figures 3 and 4). This effect was consistent with primers amplifying both the short fragment of the 12S gene, and the larger fragment of the COI gene (Figures 3 and 4). Using the 12S primer set, we found that the OTU and ZOTU richnesses recovered was consistently lower than using the COI primer set, an expected outcome due to the primer design targeting specific groups of organisms. Despite this, we also found that the analysis of five natural replicates including ten PCR replicates each, for a total of 50 PCR replicates for each mock community, we were not able to recover the whole OTU or ZOTU richness in any of the tested combinations, suggesting that a high number of natural replicate would be necessary to reach the plateau phase of the rarefaction curves (Figure 3 and 4). This was further exacerbated with the use of the primer set amplifying a larger fragment of the COI gene, which allowed the recovery of a higher ZOTU richness (Figure 4). Finally, we found that across specific habitats, the use of AMG polymerase significantly improved the OTU and ZOTU richness recovery (Figure 4) (Table S3).

**Figure 3:**
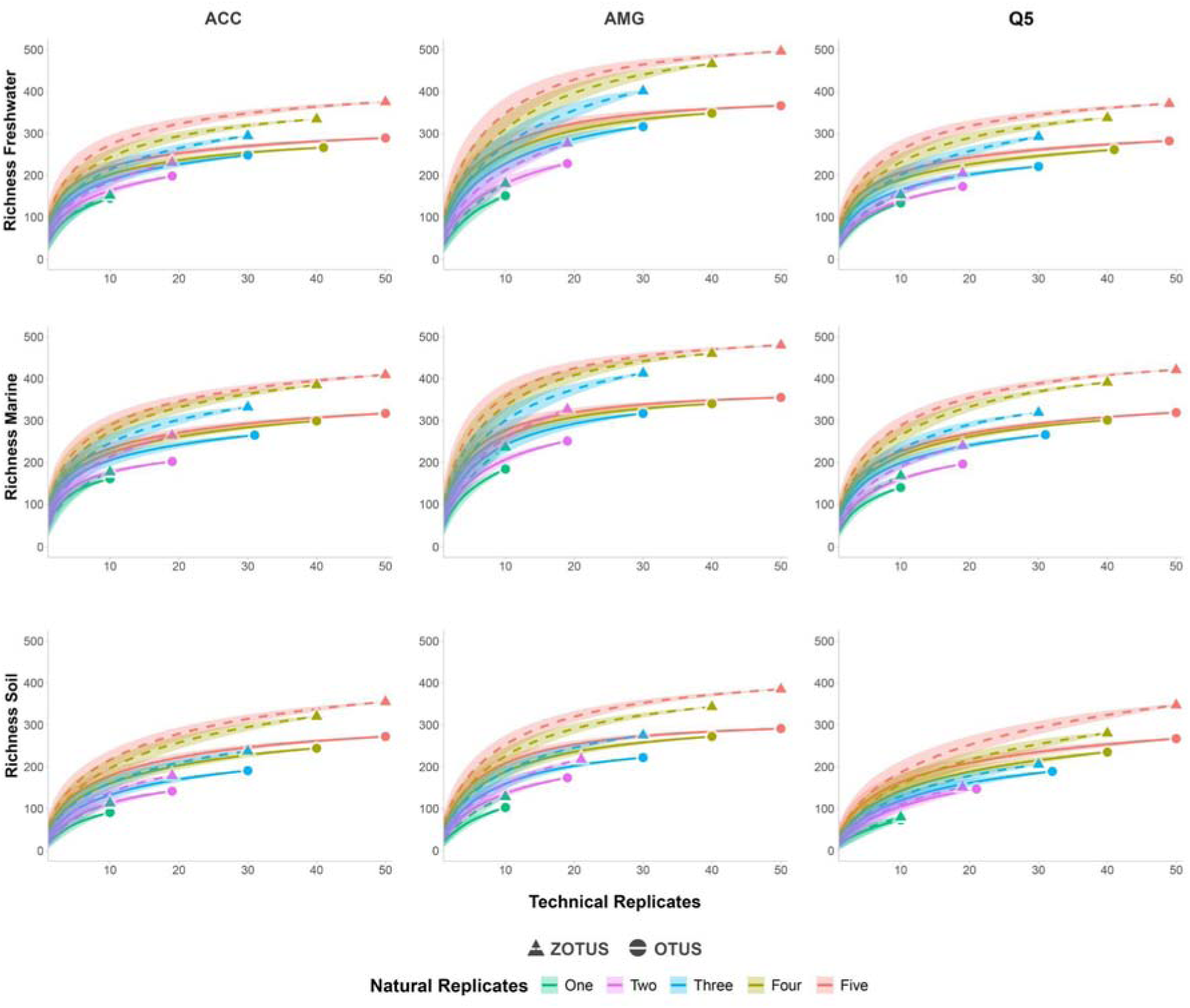
Rarefaction curves displaying the impacts of both natural replicates (i.e. number of filter or soil samples) and technical replicates (i.e. number of PCR replicates) using primers amplifying a short fragment of the 12S gene across freshwater, marine and soil habitats using Q5, ACC ang AMG polymerases. We investigated the impacts from one to five natural replicates, with up to 10 technical replicates for each natural replicates. The recovered richness represents the raw number of unique OTUs and ZOTUs detected for each combination. The shaded areas represent the confidence intervals. All graphs highlight the dissimilar impacts of natural replicates compared to technical replicates.

**Figure 4:**
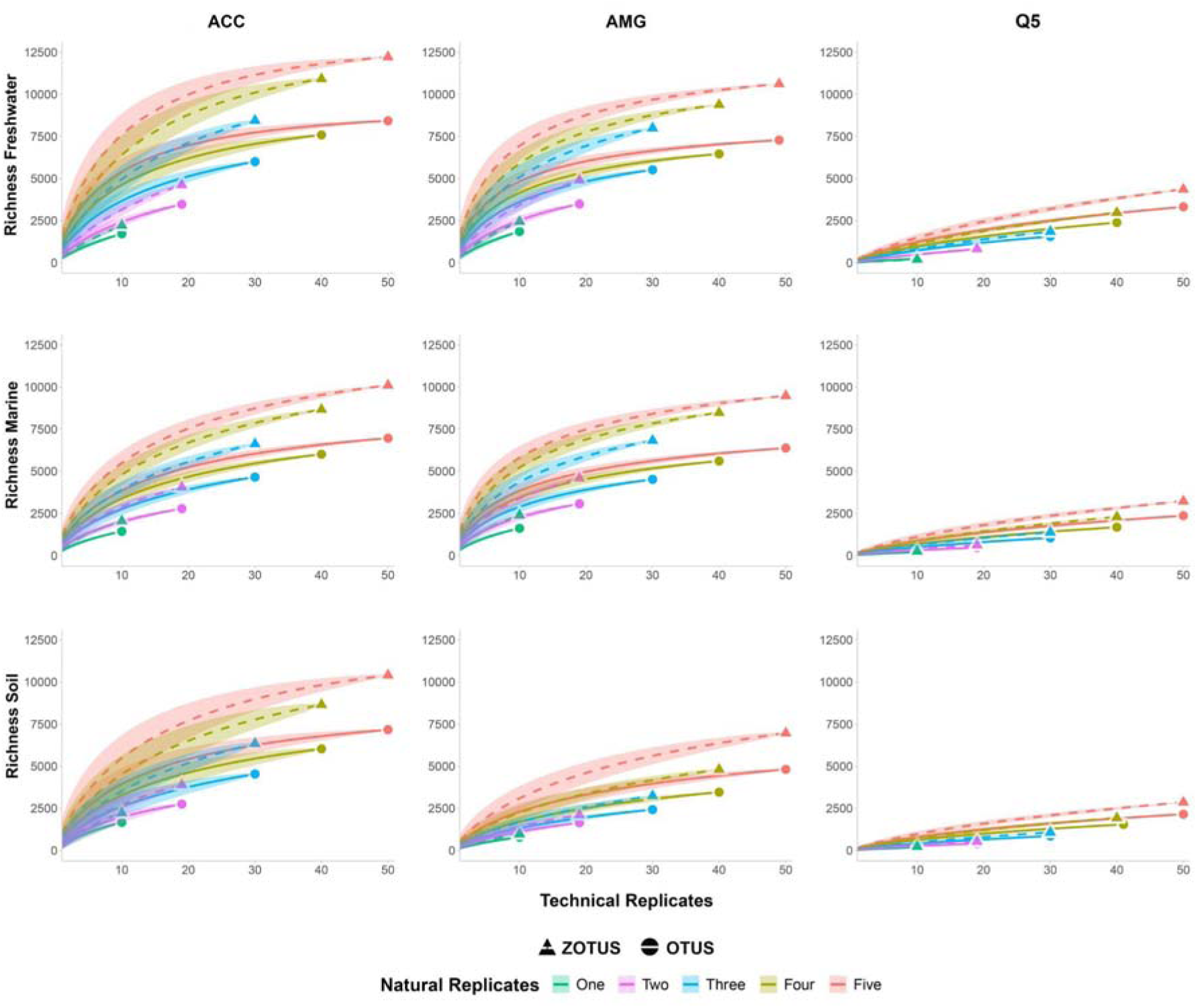
Rarefaction curves displaying the impacts of natural replicates (i.e. number of filter or soil samples) and technical replicates (i.e. number of PCR replicates) using primers amplifying a larger fragment of the COI gene across freshwater, marine and soil habitats using Q5, ACC ang AMG polymerases. As in Figure 3, we investigated the impacts from one to five natural replicates, with up to 10 technical replicates for each natural replicates. The recovered richness represents the raw number of unique OTUs and ZOTUs detected for each combination. The shaded areas represent the confidence intervals. All graphs highlight the dissimilar impacts of natural replicates compared to technical replicates, and the effect of richness recovery with each tested polymerase.

## Discussion

eDNA-based assessment is increasingly used worldwide for ecological studies and biodiversity monitoring (Thomsen & Willerslev, 2015; Taberlet et al., 2018; Deiner et al., 2021; Huang et al., 2022), and currently used in freshwater, marine, terrestrial and even air environments, allowing to quickly and efficiently monitor large areas and including citizen sciences (Agersnap & Thomsen, 2022; Bohmann & Lynggaard, 2022b; Carvalho et al., 2024). While previous work have investigated the impact of false positive and false negative (Chambert et al., 2015; Ferguson et al., 2015; Ficetola et al., 2016; Guillera-Arroita et al., 2017; Pinfield et al., 2019; Burian et al., 2021; Darling et al., 2021), payoffs of both natural and technical replicates (Ficetola et al., 2015; Beentjes et al., 2019; Mauvisseau et al., 2019; Fukaya et al., 2021), impact of primer sets on the either recovered OTU and/or ZOTU richness (Rojahn et al., 2021; Shu et al., 2021; Xiong et al., 2022; Macher et al., 2023), most studies often focus on only one set of these parameters, or on one habitat only, hampering our understanding of the combined parameters across systems.

Here, the creation of marine, freshwater and terrestrial bulk samples combined with a structured analytical design allowed us to mitigate the effects of considering results from studies that focussed only on one aspect of variation at a time. However, it should be noted that metabarcoding analysis often leads to the generation of artefact sequences, leading to OTU/ZOTU inflation. We therefore conducted our analyses using both OTU/ZOTU approaches, with and without performing the taxonomic assignment, to avoid potential biases due to artefacts as these would be excluded following taxonomic identification of the sequences. Overall, using long COI amplicons, we found no significant differences between ACC and AMG polymerases, and observed that Q5 consistently retrieved lower OTU and ZOTU richnesses. In addition to the poor amplification rates with this combination of primer and polymerase, such results could be explained by the proofreading activity displayed by Q5, leading to a decrease of amplification errors and reduction of alpha diversity (Mahé et al., 2015; Vermeulen et al., 2016; Taberlet et al., 2018). However, this observation remained consistent with and without performing the taxonomic identification of sequences, ruling out PCR errors and artefacts as a driver of these differences. Such proofreading polymerase can additionally remove mismatches at the 3’ end of primers and lead to non-specific PCR products (Mahé et al., 2015; Vermeulen et al., 2016; Taberlet et al., 2018). Such PCR products could display incorrect lengths compared to the expected amplicons, and therefore be discarded following library preparation and fragment size selection as they will be either shorter or larger than the expected amplified fragment, leading to the observed decrease of OTU/ZOTU richness. Such bottlenecks would occur prior to sequencing and bioinformatic processes, and its effect would not be able to be mitigated during the post processing steps. Here, we used a proofreading polymerase without particular modification of the PCR protocol (i.e. without the addition of three to five phosphorothioate bonds between the nucleotides and 3’ end of the primers (Mahé et al., 2015; Vermeulen et al., 2016; Taberlet et al., 2018)). This is expected to lead to an important loss of specificity due to the generation of non-specific PCR products, and potentially explain the differences observed between the different enzymes used in this study. Additionally, we didn’t observe significant differences between AMG and ACC enzymes. As ACC is designed to work with degraded samples with a high amount of inhibition, this indicates that there were no biases due to potential inhibition in our study. While we observe significant differences between Q5 and the other tested polymerases using the long amplicon size targeting the COI gene, we did not observe this using smaller amplicons targeting the 12S gene. This difference is likely due to the lower OTU/ZOTU richness obtained using the 12S primer set compared to the COI primer set (Figure 2). However, it should be noted that we didn’t formally compare results between the different primer sets tested. Indeed, these primer sets have been designed to amplify different targets (Freeland, 2016; Schenekar et al., 2020; Burian et al., 2023). As a result, any comparison of OTU/ZOTU richness retrieved with these different sets would have been inherently biassed. Future studies should therefore conduct preliminary testing using various polymerase and mock samples of their targeted environment to identify their potential limitations depending on the primer sets used. While this current study provides an extensive testing of polymerases across environments and primer sets, various other parameters, including but not limited to inhibition, acidity, salinity, temperature and volume of water sampled are also expected to impact the recovery of OTU/ZOTU richness (Bylemans et al., 2018; Deiner et al., 2018; Holman et al., 2022; Jo, 2022; Koziol et al., 2018; Pukk et al., 2021; Ruppert et al., 2019; Seymour et al., 2018). The environments tested in our study (freshwater, marine and terrestrial) have been sampled in Norway, and while we didn’t record environmental parameters susceptible to impact the recovered richness, it is expected that similar types of environments sampled elsewhere could lead to different richness recovery.

Here, we further investigated the impacts of the number of natural and technical replicates across primer sets, environments and polymerases on the recovered OTU and ZOTU richness (Figures 3, 4). We found consistent results across short and long amplicon sizes (Figure 3 and 4). Rarefaction curves using both OTUs and ZOTUs showed a steep increase of the recovered richness when increasing the number of natural replicates, and a moderate increase when increasing the number of technical replicates (Figures 3 and 4). This shows the dissimilar impacts of the replication levels, and suggests that including a higher number of natural replicates would decrease false negative detection rates. However, while the combination of five natural replicates with ten technical replicates each allowed to reach the plateau phase of richness recovery using the short 12S amplicon, this was not the case using longer COI amplicons, further highlighting discrepancy in the taxonomic coverage, and hence the sequencing depth required per sample, for these distinct primer sets. These effects were consistent across environments and polymerases despite the impacts identified earlier (Figures 3 and 4). This suggests that a higher number of natural and technical replicates would be needed to ensure an optimal diversity recovery. Previous studies have highlighted similar findings (Ficetola et al., 2015; Hinlo et al., 2017; Beentjes et al., 2019; Mauvisseau et al., 2019; Pearman et al., 2021; Shirazi et al., 2021; Stauffer et al., 2021). However, the inclusion of such a number of samples and replicates could be counter productive, as it would decrease the number of investigated locations in ecological studies due to the analytical costs. Furthermore, additional replication covering small spatial scale and temporal heterogeneity in the sampled location might have a limited effect on the overall recovery of diversity depending on the sampling design and habitat sampled. However, other factors could mitigate the negative impacts of a low replication level. In aquatic environments, the filtration of large volumes of water would allow to retrieve an increased amount of eDNA which could be diluted or have a stochastic distribution due to the species ecology (Mächler et al., 2015; Valentini et al., 2016; Peixoto et al., 2020; Wu et al., 2024). Here, we filtered one litre of water per natural replicate, and filtering a larger volume could have impacted the results of the study. However, the filtration of a larger volume of water is also associated with inhibition, potentially leading to false negatives and potential trade-offs (McKee et al., 2014; Jane et al., 2015; Goldberg et al., 2018; Uchii et al., 2019; Baudry et al., 2021; Dubreuil et al., 2021).

Finally, other tools not tested in this study, such as occupancy modelling or other analytical pipelines can also allow to mitigate potential false negative issues and increase the reliability and efficiency of eDNA based monitoring in future work (Ferguson et al., 2015; Dorazio & Erickson, 2018; Martel et al., 2020; Fukaya et al., 2021; Burian et al., 2021; Buxton et al., 2022; Burian et al., 2023). Here, we found that AMG and ACC polymerase perform equally well in marine, freshwater and terrestrial environments using longer amplicons. Based on these findings, we recommend future studies to investigate the effects of potential polymerase on the sampled environment and primers previous to the analysis of large datasets. We also found that a combination of five natural replicates with ten technical replicates each allowed a greater species richness recovery across environments. As a result, we recommend increasing the number of natural replicates when designing future sampling, potentially combined with other strategies including the filtration of larger volumes in aquatic environments, inhibitor removal and analytical frameworks when analysing the sequencing results. This does not necessarily mean that future studies should include ten technical replicates, but care should be taken to balance the level of natural and technical replication to increase results reliability within a given sequencing run. While eDNA based detection offers new opportunities for species monitoring, it should noted that this method is associated with its own benefits and biases, and care should be taken during study design to overcome potential limitations (Rees et al., 2015; Zaiko et al., 2018; Kestel et al., 2022; Couton et al., 2023).

## Supporting information

All supplementary figures, tables and files

## Acknowledgements

We would like to thank William Gromstad, Mari Engelstad and María Ariza for their valuable support with the collection of the soil and marine eDNA samples, and the Norwegian Sequencing Center for their valuable expertise.

## Author Contribution

Conceptualization: JAA, JR, HdB, QM; Sampling: JAA, ASN, QM; Laboratory analysis: JAA, LT, ASN; Bioinformatics: QM; Data curation: FdA, QM; Statistical analysis: FdA, QM, JR; Original draft: JAA, LB, ASN, QM, JR; Review and editing: JAA, LT, ASN, FdA, SMS, JR, HdB, QM; Funding acquisition: JR, HdB, QM.

## Data availability statement

Raw sequencing data can be found here: https://doi.org/10.5281/zenodo.14025716, and additional information can be found in the supplementary information.

## Declaration of competing interest

The authors declare no competing interests.

## Funding

JR, HdB and QM acknowledge funding from the Research Council of Norway through RCN INTPART 322457: SAMBA: Scaling Advanced Methods for Biodiversity Assessments; and from HK-Dir through UTF-2017-CAPES-SIU/10022: Transnational training in eDNA for biodiversity assessments and restoration ecology. We also thank Sigma2 HPC through the Biodiversity Research Consortium Brazil-Norway project METABAR NN9813K for providing computing capacities.

## Supplementary information

Supplementary Figure 1. Mean OTUs and ZOTUs numbers per treatment using the 12S primer set.

Supplementary Figure 2. Mean OTUs and ZOTUs numbers per treatment using the COI primer set.

Supplementary Figure 3. Mean raw read number per treatment using the 12S primer sets.

Supplementary Figure 4. Mean raw read number per treatment using the COI primer sets.

Supplementary Table 1. Thermal profiles used for the PCR amplifications.

Supplementary Table 2. DNA sample plate setup.

Supplementary Table 3. Pairwise comparisons of different sample types.

Supplementary Table 4. OTU table 12S

Supplementary Table 5. OTU table with taxonomic assignment 12S

Supplementary Table 6. OTU table COI

Supplementary Table 7. OTU table with taxonomic assignment COI

Supplementary Table 8. ZOTU table 12S

Supplementary Table 9. ZOTU table with taxonomic assignment 12S

Supplementary Table 10. ZOTU table COI

Supplementary Table 11. ZOTU table with taxonomic assignment COI

Supplementary File 1. Bioinformatics details and command lines.

Supplementary File 2. Custom script used for the taxonomic assignment of the COI dataset

Supplementary File 3. Custom COI reference database

## Notes

https://doi.org/10.5281/zenodo.14025716

### Competing Interest Statement

The authors have declared no competing interest.

