## Supplementary material for "eDNA replicates, polymerase and amplicon size impact inference of richness across habitats": All supplementary figures, tables and files: supplementary_figures_4.docx

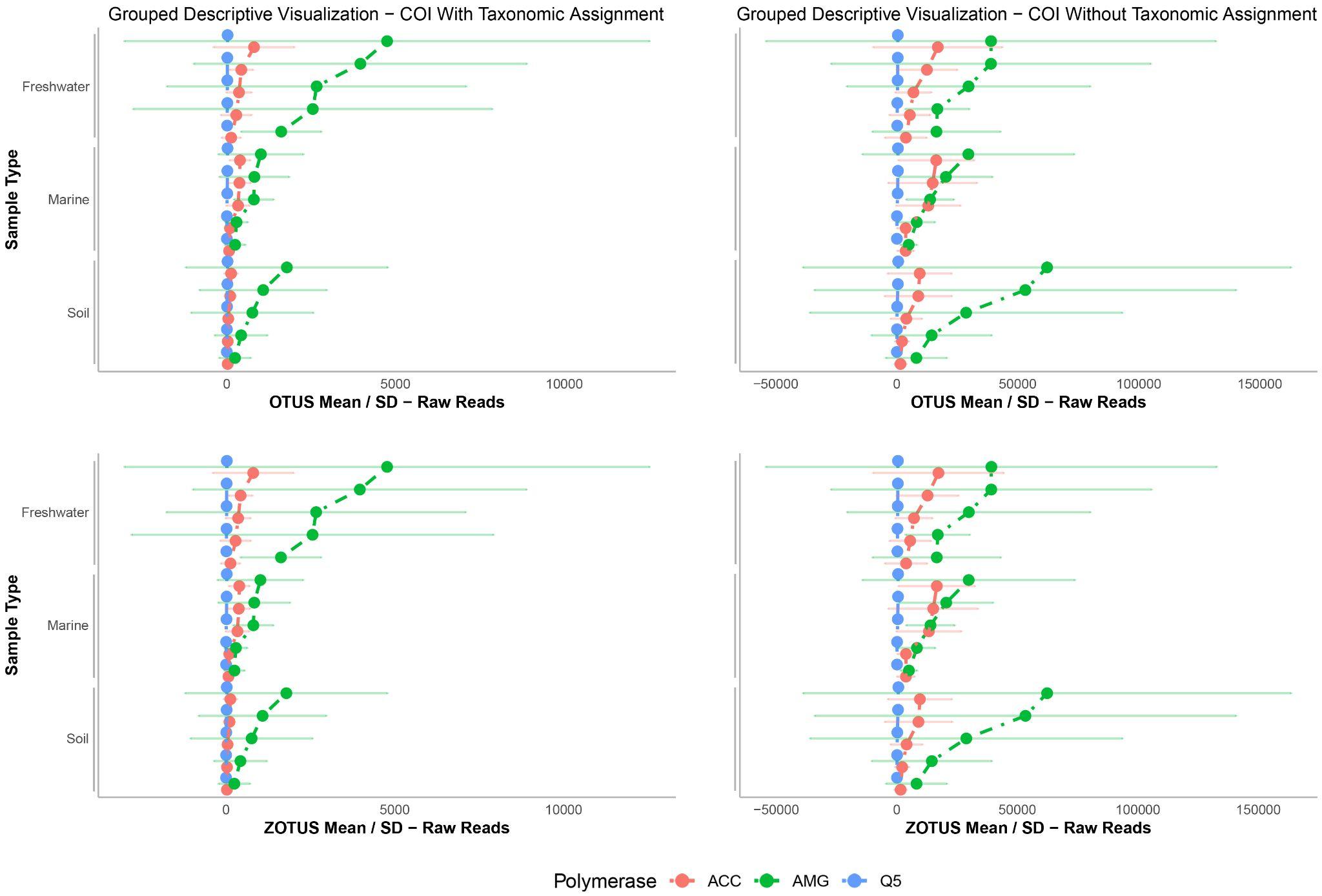

**Supplementary Figure 4**. Mean raw read number and standard deviation across the ten technical replicates per treatment (i.e. environment and polymerase) using the COI primer set, with and without taxonomic assignment.
