## Supplementary material for "eDNA replicates, polymerase and amplicon size impact inference of richness across habitats": All supplementary figures, tables and files: supplementary_file_1.docx

**Bioinformatics details and command lines**

Bioinformatics data processing, filtering and cleaning was performed as in (Carvalho et al., 2024a, b) and following the method and scripts outlined in <https://otagomohio.github.io/workshops/eDNA_Metabarcoding.html#chapter_1:_introduction> with slight modification depending on the amplicon’s libraries and regarding taxonomic assignment.

###########

The softwares necessary for the raw data processing were PEAR, OBITools, USEARCH and VSEARCH. A breakdown of the main commands used is detailed below for the analysis of the 12S library. Additional commands linked to quality control visualization are not displayed here. Similar commands using slightly different filtering parameters (highlighted in the method section) can be used to process the COI library.

1) command lines to load the necessary softwares:

module load PEAR/0.9.11-GCC-11.2.0

module load OBITools/1.2.13-GCCcore-11.2.0

module load USEARCH/11.0.667-i86linux32

module load VSEARCH/2.21.1-GCC-10.3.0

###########

Forward and reverse raw sequencing reads were merged using PEAR 0.9.3 (Zhang et al., 2014).

2) command line:

pear -f 2-12S-polproj_S1_L001_R1_001.fastq.gz -r 2-12S-polproj_S1_L001_R2_001.fastq.gz -o merged.fastq

###########

Demultiplexing was performed using the ngsfilter algorithm from OBITools (Boyer et al., 2015).

3) command line:

ngsfilter -t mapping12S.txt -u unidentified_sequences.fasta -e 2 merged.fastq > merged_demultiplexed.fastq

###########

The obigrep algorithm from OBITools was used to remove fragments >420 bp from the COI library, and >200 bp from the amplicons of the 12S library.

4) command line (for the COI library):

obigrep -L 200 merged_demultiplexed.fastq > merged_demultiplexed_L200.fastq

###########

Sample metadata were filtered out using OBITools

5) command line (for the COI library):

obiannotate -k sample merged_demultiplexed_L200.fastq > merged_demultiplexed_L200_annotated.fastq

###########

Files were then split using OBITools before further processing and place in a new folder

6) command lines:

obisplit -t sample merged_demultiplexed_L200_annotated.fastq

###########

Additional filtering was conducted using the USEARCH algorithm (Edgar, 2010, 2013, 2016), and sequences <100 bp were removed from both 12S and COI amplicon libraries.

7) command lines:

for fq in *.fastq; do usearch -fastq_filter $fq -fastq_maxee 1 -fastq_minlen 100 -fastq_maxns 0 -relabel $fq. -fastaout $fq.fasta -fastqout $fq.fastq; done

###########

Fasta files were concatenated and lower case characters were changed back to upper case characters.

8) command lines:

cat *.fasta > pooled.fasta

tr ‘[:lower:]’ ‘[:upper:]’ < pooled.fasta > pooled_upper.fasta

###########

Dereplication into unique sequences was done using USEARCH. Sequences appearing less than 10 times across the whole datasets were removed.

9) command lines:

usearch -fastx_uniques pooled_upper.fasta -fastaout uniques_10.fasta -relabel Uniq -sizeout -minuniquesize 10

###########

Finally, 12S and COI datasets were denoised using the UNOISE algorithm (Edgar, 2016) to retrieve ZOTUs (i.e. Zero Radius Operational Taxonomic Units) and clustered using the UPARSE algorithm (Edgar, 2013) to retrieve OTUs (i.e. Operational Taxonomic Units).

First, unique sequences were sorted out by abundance.

10) command lines:

module load usearch/11.0.667

usearch -sortbysize uniques_10.fasta -fastaout uniques_10_sorted.fasta

###########

Sample names were modified using:

11) command lines:

sed ’s/-/_/g’ pooled_upper.fasta > pooled_upper_changed.fasta

###########

Denoising approach:

Denoising was done using the following command:

command line:

usearch -unoise3 uniques_10_sorted.fasta -zotus zotus_10.fasta -tabbedout unoise3_10.txt

And the ZOTU table generated using the following command:

command line:

usearch -otutab pooled_upper_changed.fasta -zotus zotus_10.fasta -otutabout zotutab.txt

###########

Clustering approach:

Clustering was done using the following command:

command line:

usearch -cluster_otus uniques_10_sorted.fasta -otus otus.fasta -uparseout uparse.txt -relabel Otu

And the OTU table generated using the following command:

command line:

usearch -otutab pooled_upper_changed.fasta -otus otus.fasta -otutabout otutab.txt

###########

Taxonomic identification:

Taxonomic identification using the COI library was done using the custom script and database provided in the supplementary files.

Taxonomic identification using the 12S library was done using VSEARCH and the sintax option on the MIDORI2 database (Leray et al., 2022) (<https://www.reference-midori.info/>). Example below was done using the clustered 12S dataset.

command line:

vsearch -sintax otus.fasta -db MIDORI2_UNIQ_NUC_SP_GB255_srRNA_SINTAX_conv.udb -tabbedout OTU_12S.txt -strand both -sintax_cutoff 0.9
