## Supplementary material for "eDNA replicates, polymerase and amplicon size impact inference of richness across habitats": All supplementary figures, tables and files: supplementary_file_2.R

```
refDB_Blast <- function(Directory, Database_File, otu_table = "otu_table.txt", query = "otus.fasta", task = "megablast", out = "blast.txt", 
                        max_target_seqs = 50, perc_identity = 95, qcov_hsp_perc = 95, num_threads = 6, 
                        Specie_Threshold = 99, Genus_Threshold = 97, Family_Threshold = 95, 
                        penalty = NULL, reward = NULL, evalue = NULL, word_size = NULL, gapopen = NULL, 
                        gapextend = NULL, max_hsps = NULL, xdrop_ungap = NULL, xdrop_gap = NULL, 
                        xdrop_gap_final = NULL, searchsp = NULL, sum_stats = NULL, no_greedy = NULL, 
                        min_raw_gapped_score = NULL, template_type = NULL, template_length = NULL, 
                        dust = NULL, filtering_db = NULL, window_masker_taxid = NULL, window_masker_db = NULL, 
                        soft_masking = NULL, ungapped = NULL, culling_limit = NULL, 
                        best_hit_overhang = NULL, best_hit_score_edge = NULL, subject_besthit = NULL, 
                        window_size = NULL, off_diagonal_range = NULL, use_index = NULL, index_name = NULL, 
                        lcase_masking = NULL, query_loc = NULL, strand = NULL, parse_deflines = NULL, 
                        outfmt = "6", show_gis = NULL, num_descriptions = NULL, num_alignments = NULL, 
                        line_length = NULL, html = NULL, sorthits = NULL, sorthsps = NULL, 
                        mt_mode = NULL, remote = NULL) {
  
  linux_path <- gsub("C:/", "/mnt/c/", Directory)
  
  # Optional Parameters
  optional_params <- ""
  add_param <- function(param, value) {
    if (!is.null(value)) {
      return(paste0(" -", param, " ", value))
    }
    return("")
  }
  
  optional_params <- paste0(optional_params, 
                            add_param("penalty", penalty), 
                            add_param("reward", reward), 
                            add_param("evalue", evalue),
                            add_param("word_size", word_size),
                            add_param("gapopen", gapopen),
                            add_param("gapextend", gapextend),
                            add_param("max_hsps", max_hsps),
                            add_param("xdrop_ungap", xdrop_ungap),
                            add_param("xdrop_gap", xdrop_gap),
                            add_param("xdrop_gap_final", xdrop_gap_final),
                            add_param("searchsp", searchsp),
                            add_param("sum_stats", sum_stats),
                            add_param("no_greedy", no_greedy),
                            add_param("min_raw_gapped_score", min_raw_gapped_score),
                            add_param("template_type", template_type),
                            add_param("template_length", template_length),
                            add_param("dust", dust),
                            add_param("filtering_db", filtering_db),
                            add_param("window_masker_taxid", window_masker_taxid),
                            add_param("window_masker_db", window_masker_db),
                            add_param("soft_masking", soft_masking),
                            add_param("ungapped", ungapped),
                            add_param("culling_limit", culling_limit),
                            add_param("best_hit_overhang", best_hit_overhang),
                            add_param("best_hit_score_edge", best_hit_score_edge),
                            add_param("subject_besthit", subject_besthit),
                            add_param("window_size", window_size),
                            add_param("off_diagonal_range", off_diagonal_range),
                            add_param("use_index", use_index),
                            add_param("index_name", index_name),
                            add_param("lcase_masking", lcase_masking),
                            add_param("query_loc", query_loc),
                            add_param("strand", strand),
                            add_param("parse_deflines", parse_deflines),
                            add_param("outfmt", outfmt),
                            add_param("show_gis", show_gis),
                            add_param("num_descriptions", num_descriptions),
                            add_param("num_alignments", num_alignments),
                            add_param("line_length", line_length),
                            add_param("html", html),
                            add_param("sorthits", sorthits),
                            add_param("sorthsps", sorthsps),
                            add_param("mt_mode", mt_mode),
                            add_param("remote", remote)
  )
  
  # blastn
  system(paste0("wsl blastn -query ", paste0(linux_path, query), " -task ", task, " -db ", paste0(linux_path, Database_File), 
                " -out ", out, " -max_target_seqs ", max_target_seqs, 
                " -perc_identity ", perc_identity, 
                " -qcov_hsp_perc ", qcov_hsp_perc, 
                " -num_threads ", num_threads, 
                optional_params))
  
  system(paste0("wsl blastn -query ", paste0(linux_path, query), 
                " -task ", task, 
                " -db ", paste0(linux_path, Database_File), 
                " -out ", out, 
                " -outfmt 6",  # Set output format to tabular
                " -max_target_seqs 50", 
                " -perc_identity 95", 
                " -qcov_hsp_perc 95", 
                " -num_threads 6"))
  
  system(paste0("wsl tr -d '#' < ", otu_table, " > temp_raw_database && mv temp_raw_database ", otu_table))
  
  csv1 <- read.table(paste0(Directory, out), sep = "", header = FALSE)
  
  Samples <- read.table(paste0(Directory, otu_table), sep = "\t", header = TRUE)
  
  colnames(Samples)[1] <- "qseqid"
  
  ########## Files = csv + ID taxa ##########
  colnames(csv1) <- c("qseqid", "seqid", "pident", "length", "mismatch", "gapopen", "qstart", "qend", "sstart", "send", "evalue", "bitscore")
  
  # Separate only the max value of match per ZOTU
  cl.max_ <- csv1 %>% group_by(qseqid) %>% top_n(1, pident)
  
  # Organizing the species names
  cl.max_$seqid <- gsub("\\d+_", "", cl.max_$seqid)
  cl.max_$seqid <- gsub("_", " ", cl.max_$seqid)
  cl.max_$seqid <- sub("\\b(sp\\.?\\s+).*", "\\1", cl.max_$seqid)
  
  # Split each cell into "characters" separated by blank spaces
  split_seqid <- strsplit(cl.max_$seqid, "\\s+")
  
  # Exclude characters after the second "character" of each cell
  cl.max_$seqid <- sapply(split_seqid, function(chars) {
    paste(chars[1:2], collapse = " ")
  })
  
  # Separate only the min value of match per OTU - Evalue 
  cl.max_ <- cl.max_ %>% group_by(qseqid) %>% top_n(-1, evalue)
  
  # Separate only the min value of match per OTU - Bitscore 
  cl.max_ <- cl.max_ %>% group_by(qseqid) %>% top_n(-1, bitscore)
  
  cl.max_ <- cl.max_[!duplicated(cl.max_[1]), ]
  
  ids <- paste(cl.max_$qseqid, cl.max_$seqid, sep = " ")
  
  # data source: NCBI
  db_download_ncbi()
  src <- src_ncbi()
  
  taxa_to_id <- name2taxid(sapply(strsplit(ids, " "), function(x) paste(x[2:3], collapse = " ")))
  
  taxa_names <- classification(taxa_to_id, db = "ncbi")
  
  # Function to check if an object is a DataFrame
  is_dataframe <- function(obj) {
    inherits(obj, "data.frame")
  }
  
  # Iterate through each sublist
  for (i in seq_along(taxa_names)) {
    # If it's not a DataFrame, create a new DataFrame with NA values
    if (!is_dataframe(taxa_names[[i]])) {
      taxa_names[[i]] <- data.frame(name = NA, rank = NA, id = NA)
    }
  }
  
  # Replace names based on the vector
  names(taxa_names) <- ids
  
  taxa_names_df <- do.call(rbind, lapply(names(taxa_names), function(name) {
    df <- taxa_names[[name]]
    df$Taxa <- name
    return(df)
  }))
  
  sum(grepl("phylum", taxa_names_df$rank, ignore.case = TRUE))
  
  taxa_names_df <- taxa_names_df[, c("Taxa", names(taxa_names_df)[1:(ncol(taxa_names_df) - 1)])] 
  taxa_names_df <- taxa_names_df[, -ncol(taxa_names_df)]
  
  taxa_names_df <- taxa_names_df %>%
    filter(rank %in% c("kingdom", "phylum", "class", "order", "family", "genus", "species", NA)) %>%
    group_by(Taxa) %>%
    distinct()
  
  taxa_remaining <- taxa_names_df$Taxa[is.na(taxa_names_df$rank)]
  
  # Check if taxa_remaining is not empty
  if (length(taxa_remaining) > 0) {
    # If taxa_remaining is not empty, run this block
    taxa_with_na <- sapply(strsplit(taxa_remaining, " "), "[", 2)
    taxa_with_na <- data.frame(name = taxa_with_na)
    row.names(taxa_with_na) <- taxa_remaining
    
    # Add the ID column to taxa_with_na based on matching names
    taxa_with_na_rank <- name2taxid(taxa_with_na$name, out_type = "summary")
    taxa_with_na_with_id <- left_join(taxa_with_na, taxa_with_na_rank, by = "name")
    row.names(taxa_with_na_with_id) <- taxa_remaining
    
    # Match the values in taxa with the values in df$name
    remaining_taxa <- classification(taxa_with_na_with_id$id, db = "ncbi")
    names(remaining_taxa) <- taxa_remaining
    
    # Iterate through each sublist
    for (i in seq_along(remaining_taxa)) {
      # If it's not a DataFrame, create a new DataFrame with NA values
      if (!is_dataframe(remaining_taxa[[i]])) {
        remaining_taxa[[i]] <- data.frame(name = NA, rank = NA, id = NA)
      }
    }
    
    taxa_names_df_rem <- do.call(rbind, lapply(names(remaining_taxa), function(name) {
      df <- remaining_taxa[[name]]
      df$Taxa <- name
      return(df)
    }))
    
    taxa_names_df_rem <- taxa_names_df_rem[, c("Taxa", names(taxa_names_df_rem)[1:(ncol(taxa_names_df_rem) - 1)])]
    taxa_names_df_rem <- taxa_names_df_rem[, -ncol(taxa_names_df_rem)]
    
    taxa_names_df <- taxa_names_df %>%
      filter(rank %in% c("kingdom", "phylum", "class", "order", "family", "genus", "species", NA)) %>%
      group_by(Taxa) %>%
      distinct()
    
    taxa_names_df <- taxa_names_df[complete.cases(taxa_names_df), ]
    
    # Combine the dataframes using rbind
    all_taxa <- data.frame(rbind(taxa_names_df, taxa_names_df_rem))
    
    identified_otus <- all_taxa %>% 
      pivot_wider(names_from = rank, values_from = name)
    
  } else {
    # If taxa_remaining is empty, run this block
    taxa_names_df <- taxa_names_df[complete.cases(taxa_names_df), ]
    
    identified_otus <- taxa_names_df %>%
      pivot_wider(names_from = rank, values_from = name)
  }
  
  # Split the content based on the first blank space
  split_content <- str_split(identified_otus$Taxa, " ", n = 2)
  
  # Extract the first part (characters before the first blank space)
  first_part <- sapply(split_content, function(x) x[1])
  
  # Extract the second part (characters after the first blank space)
  second_part <- sapply(split_content, function(x) ifelse(length(x) > 1, x[2], ""))
  
  # Create new columns in the dataframe for the split content
  identified_otus <- cbind(qseqid = first_part, seqid = second_part, identified_otus[2:ncol(identified_otus)])
  
  colnames(identified_otus) <- c("qseqid", "seqid", "Kingdom", "Phylum", "Class", "Order", "Family", "Genus", "Species")
  
  cl.max_subset <- cl.max_[, c(1, 3)]
  
  cl.max_ordered <- cl.max_subset[match(identified_otus$qseqid, cl.max_subset$qseqid), ]
  
  cl.max <- cbind(identified_otus[, 1:9], cl.max_ordered$pident)
  
  colnames(cl.max)[ncol(cl.max)] <- "pident"
  
  ########## File = Samples ##########
  
  Samples <- Samples[order(match(Samples$qseqid, cl.max$qseqid)), ]
  
  Final_Match_samples <- plyr::match_df(Samples, cl.max, on = "qseqid")
  
  Blast <- cbind(cl.max, Final_Match_samples[, 2:length(colnames(Final_Match_samples))])
  
  Blast$Species <- ifelse(Blast$pident >= Genus_Threshold & Blast$pident < Specie_Threshold, "" ,Blast$Species)
  Blast$Species <- ifelse(Blast$pident >= Family_Threshold & Blast$pident < Genus_Threshold, "" ,Blast$Species)
  Blast$Genus <- ifelse(Blast$pident >= Family_Threshold & Blast$pident < Genus_Threshold, "" ,Blast$Genus)
  
  list_columns <- sapply(Blast, is.list)
  list_columns_names <- names(list_columns[list_columns])
  
  Blast[list_columns_names] <- lapply(Blast[list_columns_names], function(col) {
    sapply(col, function(x) {
      if (is.null(x)) {
        return(NA)
      } else if (is.list(x) || length(x) > 1) {
        return(as.character(x[[1]]))  
      } else {
        return(as.character(x)) 
      }
    })
  })
  
  Blast <- as.data.frame(Blast, stringsAsFactors = FALSE)
  
  write.table(Blast, file = paste0(Directory, "taxonomic_assignment.txt"), sep = "\t", row.names = F, col.names = T)
  
  # Find the Zotus not identified with perc_identity = 97
  remaining_sequences <- subset(Samples$qseqid, !(Samples$qseqid %in% unique(csv1$qseqid)))
  
  if (length(remaining_sequences) != 0) { 
    # Create a file with a list of Zotus not identified to first threshold of perc_identity
    write.table(remaining_sequences, file = paste0(Directory, "remaining_sequences.txt"), sep = " ", row.names = F, col.names = F, quote = F)
    print("End of Run")
  } else {
    print("End of Run")
  }
}
```
