## Supplementary material for "eDNA replicates, polymerase and amplicon size impact inference of richness across habitats": All supplementary figures, tables and files: supplementary_tables_1.docx

|  | Enzyme | Q5 | Amplitaq | Accustart | Accustart |
| --- | --- | --- | --- | --- | --- |
|  | Marker | *COI*/*12S* | *COI/12S* | *COI* | *12S* |
| Initial denaturation | Temp (°C) | 98 | 95 | 94 | 94 |
|  | Time | 1 min | 5 min | 2 min | 2 min |
| Denaturation | Temp (°C) | 98 | 95 | 94 | 94 |
|  | Time | 10 sec | 30 sec | 10 sec | 10 sec |
| Annealing | Temp (°C) | 59 | 45 | 47 | 55 |
|  | Time | 20 s | 30 sec | 20 sec | 20 sec |
| Elongation^a^ | Temp (°C) | 72 | 72 | 72 | 72 |
|  | Time | 10 sec | 20 sec | 15 sec | 15 sec |
| Cycles |  | 35 | 35 | 30 | 31 |

^a^A final elongation step of 2 minutes was added to all the thermal cycling programs.

**Supplementary table 1.** Thermal profiles used for the PCR amplification using the different polymerases and primers combinations.
