## Supplementary material for "eDNA replicates, polymerase and amplicon size impact inference of richness across habitats": All supplementary figures, tables and files: supplementary_tables_2.docx

|  | **1** | **2** | **3** | **4** | **5** | **6** | **7** | **8** | **9** | **10** | **11** | **12** |
| --- | --- | --- | --- | --- | --- | --- | --- | --- | --- | --- | --- | --- |
| **A** | S1-COI-Q5 | S9-COI-Q5 | S1-COI-AMG | S9-COI-AMG | S1-COI-ACC | S9-COI-ACC | S1-12S-Q5 | S9-12S-Q5 | S1-12S-AMG | S9-12S-AMG | S1-12S-ACC | S9-12S-ACC |
| **B** | S2-COI-Q5 | S10-COI-Q5 | S2-COI-AMG | S10-COI-AMG | S2-COI-ACC | S10-COI-ACC | S2-12S-Q5 | S10-12S-Q5 | S2-12S-AMG | S10-12S-AMG | S2-12S-ACC | S10-12S-ACC |
| **C** | S3-COI-Q5 | S11-COI-Q5 | S3-COI-AMG | S11-COI-AMG | S3-COI-ACC | S11-COI-ACC | S3-12S-Q5 | S11-12S-Q5 | S3-12S-AMG | S11-12S-AMG | S3-12S-ACC | S11-12S-ACC |
| **D** | S4-COI-Q5 | S12-COI-Q5 | S4-COI-AMG | S12-COI-AMG | S4-COI-ACC | S12-COI-ACC | S4-12S-Q5 | S12-12S-Q5 | S4-12S-AMG | S12-12S-AMG | S4-12S-ACC | S12-12S-ACC |
| **E** | S5-COI-Q5 | S13-COI-Q5 | S5-COI-AMG | S13-COI-AMG | S5-COI-ACC | S13-COI-ACC | S5-12S-Q5 | S13-12S-Q5 | S5-12S-AMG | S13-12S-AMG | S5-12S-ACC | S13-12S-ACC |
| **F** | S6-COI-Q5 | S14-COI-Q5 | S6-COI-AMG | S14-COI-AMG | S6-COI-ACC | S14-COI-ACC | S6-12S-Q5 | S14-12S-Q5 | S6-12S-AMG | S14-12S-AMG | S6-12S-ACC | S14-12S-ACC |
| **G** | S7-COI-Q5 | S15-COI-Q5 | S7-COI-AMG | S15-COI-AMG | S7-COI-ACC | S15-COI-ACC | S7-12S-Q5 | S15-12S-Q5 | S7-12S-AMG | S15-12S-AMG | S7-12S-ACC | S15-12S-ACC |
| **H** | S8-COI-Q5 | NTC-COI-Q5 | S8-COI-ANG | NTC-COI-AMG | S8-COI-ACC | NTC-COI-ACC | S8-12S-Q5 | NTC-12S-Q5 | S8-12S-AMG | NTC-12S-AMG | S8-12S-ACC | NTC-12S-ACC |

**Supplementary table 2**: Sample plate setup used in this study. This plate design was replicated in ten independent PCR setups. Yielding a total of 960 amplicons with both primer sets combined. The samples S1-S5 are from the marine environment, S6-S10 are from the limnic environment and S11-S15 are the soil samples. COI/12S are the two markers amplified (using the Leray and Riaz primer sets) and Q5/AMG/ACC are the three enzymes Q5 polymerase, Amplitaq Gold 360 and Accustart Toughmix, respectively. The NTCs are non-template controls (dH_2_O instead of biological sample).
