## Supplementary material for "eDNA replicates, polymerase and amplicon size impact inference of richness across habitats": All supplementary figures, tables and files: supplementary_tables_3.docx

| **Marker** | **Approach** | **Sample Type** | **Test** | **Pairwise Comparison** | **p-value** |
| --- | --- | --- | --- | --- | --- |
| COI | OTU | Freshwater | Kruskal-Wallis | - | **0.000** |
| COI | OTU | Freshwater | Dunn's Test | ACC - AMG | 0.665 |
| COI | OTU | Freshwater | Dunn's Test | ACC - Q5 | **0.000** |
| COI | OTU | Freshwater | Dunn's Test | AMG - Q5 | **0.000** |
| COI | OTU | Marine | Kruskal-Wallis | - | **0.000** |
| COI | OTU | Marine | Dunn's Test | ACC - AMG | 0.361 |
| COI | OTU | Marine | Dunn's Test | ACC - Q5 | **0.000** |
| COI | OTU | Marine | Dunn's Test | AMG - Q5 | **0.000** |
| COI | OTU | Soil | Kruskal-Wallis | - | **0.000** |
| COI | OTU | Soil | Dunn's Test | ACC - AMG | **0.047** |
| COI | OTU | Soil | Dunn's Test | ACC - Q5 | **0.000** |
| COI | OTU | Soil | Dunn's Test | AMG - Q5 | **0.000** |
| COI | ZOTU | Freshwater | Kruskal-Wallis | - | **0.000** |
| COI | ZOTU | Freshwater | Dunn's Test | ACC - AMG | 0.554 |
| COI | ZOTU | Freshwater | Dunn's Test | ACC - Q5 | **0.000** |
| COI | ZOTU | Freshwater | Dunn's Test | AMG - Q5 | **0.000** |
| COI | ZOTU | Marine | Kruskal-Wallis | - | **0.000** |
| COI | ZOTU | Marine | Dunn's Test | ACC - AMG | 0.361 |
| COI | ZOTU | Marine | Dunn's Test | ACC - Q5 | **0.000** |
| COI | ZOTU | Marine | Dunn's Test | AMG - Q5 | **0.000** |
| COI | ZOTU | Soil | Kruskal-Wallis | - | **0.000** |
| COI | ZOTU | Soil | Dunn's Test | ACC - AMG | **0.048** |
| COI | ZOTU | Soil | Dunn's Test | ACC - Q5 | **0.000** |
| COI | ZOTU | Soil | Dunn's Test | AMG - Q5 | **0.000** |
| 12s | OTU | Freshwater | Kruskal-Wallis | - | 0.444 |
| 12s | OTU | Freshwater | Dunn's Test | ACC - AMG | 0.978 |
| 12s | OTU | Freshwater | Dunn's Test | ACC - Q5 | 0.313 |
| 12s | OTU | Freshwater | Dunn's Test | AMG - Q5 | 0.628 |
| 12s | OTU | Marine | Kruskal-Wallis | - | 0.324 |
| 12s | OTU | Marine | Dunn's Test | ACC - AMG | 0.331 |
| 12s | OTU | Marine | Dunn's Test | ACC - Q5 | 0.260 |
| 12s | OTU | Marine | Dunn's Test | AMG - Q5 | 0.999 |
| 12s | OTU | Soil | Kruskal-Wallis | - | 0.103 |
| 12s | OTU | Soil | Dunn's Test | ACC - AMG | 0.999 |
| 12s | OTU | Soil | Dunn's Test | ACC - Q5 | 0.143 |
| 12s | OTU | Soil | Dunn's Test | AMG - Q5 | 0.070 |
| 12s | ZOTU | Freshwater | Kruskal-Wallis | - | 0.522 |
| 12s | ZOTU | Freshwater | Dunn's Test | ACC - AMG | 0.999 |
| 12s | ZOTU | Freshwater | Dunn's Test | ACC - Q5 | 0.439 |
| 12s | ZOTU | Freshwater | Dunn's Test | AMG - Q5 | 0.546 |
| 12s | ZOTU | Marine | Kruskal-Wallis | - | 0.727 |
| 12s | ZOTU | Marine | Dunn's Test | ACC - AMG | 0.901 |
| 12s | ZOTU | Marine | Dunn's Test | ACC - Q5 | 0.650 |
| 12s | ZOTU | Marine | Dunn's Test | AMG - Q5 | 0.999 |
| 12s | ZOTU | Soil | Kruskal-Wallis | - | 0.140 |
| 12s | ZOTU | Soil | Dunn's Test | ACC - AMG | 0.999 |
| 12s | ZOTU | Soil | Dunn's Test | ACC - Q5 | 0.280 |
| 12s | ZOTU | Soil | Dunn's Test | AMG - Q5 | 0.110 |

**Supplementary table S3.** Pairwise comparisons of different sample types (Freshwater, Marine, and Soil) for COI and 12S markers using OTU and ZOTU approaches. The Kruskal-Wallis test was used to assess the overall differences, and Dunn's Test was applied for pairwise comparisons between groups: ACC, AMG, and Q5. The p-values are reported for each comparison, with significant p-values (p < 0.05) indicating statistically significant differences between the groups. A p-value of 0.000 indicates values less than 0.001.
